# Constraining Palaeogeography and Palaeotides for the Cambrian using cnidarian medusae

**DOI:** 10.64898/2026.08.27.747545

**Authors:** H. A. M. Byrne, M. E. H. Hartley, I. Perez, C. R. Scotese, D. J. Lunt, P. J. Valdes, J. A. M. Green

## Abstract

The ocean tides influence key Earth system processes at a range of spatial and temporal scales. It is known that the geometry of ocean basins is the leading controller of tidal energetics, so well-constrained palaeogeographic reconstructions and tidal properties for Earth’s past are imperative when investigating other Earth system processes. Here, we present a novel way to constrain both deep-time tidal model results and reconstructions, by combining palaeoecology with sedimentology. We compare new palaeo-tidal model simulations for the Cambrian period, significant for the early origin and radiation of major animal fauna, to tidal proxies. One of the most abundant soft-bodied organisms preserved during this time are cnidarian medusae (“jellyfish”). A total of 17 cnidarian medusae localities were obtained through the literature, which had an adequate global distribution and occurred at regular intervals throughout the period of study. In some locations there were also estimates of palaeo-tidal range. Our results show a good agreement between the simulations and proxy data. In the few locations where there is disagreement, it is proposed that the palaeogeographic reconstructions are missing details, e.g., island chains, and our results allow for the palaeogeographic reconstructions to be improved. The proxy method presented is promising and can be applied to other time-periods with different marine fossils, particularly at evolutionary and extinction periods where the marginal marine environment is of importance.

**Key Points:**

- First use of palaeoecology combined with sedimentology to constrain both reconstructions and tidal simulation outputs, trialled in this study on the Cambrian period.
- Good agreement between proxy and simulation data for tidal ranges.
- Proxy method holds potential for use with different marine fossils to examine other time periods.

**Plain Language Summary:** Ocean tides are crucial for shaping Earth’s environment on short and long timescales. The shape and layout of ocean basins are the main factors that influence how tides behave. To understand how tides affected Earth’s history, we need accurate models of ancient ocean conditions and how they interacted with the environment. Here, we use a new approach to improve tidal models by combining fossil evidence with sediment analysis. We focused on the Cambrian period, a time when many major animal groups first evolved. One of the most common fossils from this time are jellyfish. We investigated 17 locations where jellyfish fossils have been found by looking at the surrounding sediments for signs of tidal influences, including tidal range. Our tidal models generally matched the data from the fossil sites, but where they didn’t, we believe it’s because the ancient maps are missing features, like small islands. This can be used for other time periods, especially those tied to important evolutionary or extinction events in shallow marine environments. We believe there is poetry in this type of research and provide an example in the supplementary material.

## 1. Introduction

Palaeogeographic reconstructions are paramount for deep time-research, e.g., when exploring biogeographical links between fossil groups, and to simulate past climate systems (De Vleeschouwer et al., 2014; Dowding & Ebach, 2019; Farnsworth et al., 2019; Laugié et al., 2020; Valdes et al., 2021; S. -B. Wilmes et al., 2021; Biewald et al, 2024). Recent advances in methods mean that reconstructions can now cover 1.8 billion years, from the Neoproterozoic to the present, and a further 200-250 million years (My) into the future (Cao et al. 2017; Davies et al., 2018; Mather et al., 2024; Merdith et al., 2021; Scotese, 2021; Scotese & Wright, 2021). Furthermore, the latest updates to the reconstructions now include information about land topography and ocean depths for the Phanerozoic (i.e., 540 Ma-present; note that Ma is million years ago in the following). Given the essential role palaeogeographic reconstructions play in deep-time research, it is imperative to constrain them, particularly with regards to the bathymetry which is difficult to determine as the vast majority of present-day oceanic crust is younger than 280 Ma (Müller et al., 2008).

The ocean tide is an important process in the Earth system, and having well-constrained estimates of tidal properties for Earth’s past is crucial when investigating other Earth system processes, including ocean circulation (Green & Huber, 2013; Ladant et al., 2024; Wilmes et al., 2021; Wunsch & Ferrari, 2004), biogeochemistry (Sharples et al., 2007; Wilmes et al., 2021), sedimentology (Davis & Dalrymple, 2011), climate (Ladant et al., 2024; Laugié et al., 2020), and the evolution of the Earth’s orbit (e.g., Daher et al., 2021). Deep-time tides are usually explored using numerical tidal models, which require validation using some form of tidal proxy (Byrne et al., 2020; Green et al., 2020; Guo et al., 2023; Zuchuat et al., 2022). To date, tidal proxy information has mainly come from tidalites, because their alternating layers of sand and mud are interpreted as being deposited by the ebb-flood tidal cycle (Klein, 1971; Longhitano et al., 2012; Visser, 1980). However, there are concerns that the layers may not be complete or laid down by other cyclic processes, e.g., non-tidal estuarine and lacustrine conditions, potentially leading to an overinterpretation of the information provided (Dalrymple, 2021; Gugliotta et al., 2016; Tessier, 1998). Consequently, there is a need for alternative proxies for tides. At the same time, tides are to first order controlled by ocean basin geometry (Blackledge et al., 2020; J. A. M. Green et al., 2017, 2018; M. Green et al., 2023), so if we can constrain that, e.g., by identifying the location of coastlines, we can be confident in our tidal model simulations. To address the need for more robust proxies to constrain both reconstructions and tidal simulations at once, we propose the use of palaeoecology in conjunction with sedimentology. Palaeoecology can be used as a powerful palaeoenvironment indicator and has been successfully used to constrain reconstructions during the later Palaeozoic and the palaeogeographic history of Australia (Cao et al., 2017; Kocsis & Scotese, 2021; Wright et al., 2013). If paleoecology gives us confidence that a deposit is marine, the geology itself can occasionally be useful if an estimate of the tidal range can be obtained (Guo et al., 2023). These metrics can then be compared to tidal simulation outputs as a validation. This is the first study to use this two-fold proxy approach - combining palaeoecology and sedimentology – to explicitly constrain tidal model simulations, and it will be tested on the earliest reconstructions with an abundant fossil record: the Cambrian Period.

The Cambrian Period lasted between 538.8-485.4 Ma and was home to pronounced changes in life on Earth. More complex multicellular organisms had emerged during the preceding Ediacaran (Butterfield, 2007), but mineralised organisms became common during the Cambrian, and the rapid diversification in the Cambrian explosion led to the first representation of all present-day animal phyla (Benton & Harper, 2020). The Cambrian explosion has been spectacularly captured in an abundant number of localities of exceptional preservation, known as Lagerstätten (Wood et al., 2019). Interestingly, the Cambrian contains a high proportion of Lagerstätten critically preserving many soft-bodied organisms, and so our understanding of the Cambrian biology surpasses that of some later periods (Orr et al., 2003). Among the evolutionary highlights of the Cambrian are the rise of predators, instigating an arms race between predator apparatus and prey defence, and faunal disparity (Porter, 2011), with a shift in biota from shelf seas into shallower waters (Wood et al., 2019).

The phylum Cnidaria, which includes corals and medusozoans (jellyfish), has an extensive spatial and temporal fossil record from the Cambrian period. Fossilised Cnidarians from the period are found in a broad range of marine settings, predominantly in the intertidal zone, making them an ideal group for palaeoecological study (Young & Hagadorn, 2020). Here, we focused on fossil medusozoans and extracted the palaeoenvironment inferences from their palaeoecology. The resulting information is then compared to simulations using an established tidal model. The boundary conditions for the model are palaeogeographic reconstructions from Wright & Scotese (2018) at 5 Ma intervals throughout the Cambrian, beginning at 540 Ma (terminal Ediacaran) to 485 Ma (terminal Cambrian-earliest Ordovician). Major modifications to the reconstructions based on our results are beyond the scope of this study, but minor alterations to bathymetries will be done and will allow for reconstruction alterations to be made in future research.

## 2 Methods

### 2.1 Palaeogeographic reconstructions

The Cambrian explosion occurred against the backdrop of a very different looking earth to present day, with a configuration consisting of four main continents, mostly concentrated in the southern hemisphere; Baltica, Gondwana, Laurentia, and Siberia (see Fig. 1a). The palaeo-bathymetries used here came from the Digital Elevation Models (DEMs) presented by Scotese and Wright (2018) and described in more detail in Scotese (2021). The data was supplied at 1/10° horizontal resolution for every 5 Myr. Shelf seas in the DEM were ∼170 m deep all the way to the coast. Consequently, to obtain a more realistic bathymetry, we ran a 5-point moving average in both latitude and longitude. The size of the averaging was chosen because it removes small-scale coastal variability present in the paleoDEMs, and whilst this means the zonal length of the smoothing varies with latitude there was little discernible difference when compared to one DEM where the averaging was done over distance. All bathymetries effectively ran from 89°S to 89°N in latitude because of the introduction of land in the tidal model covering the poles due to the convergence of the grid cells there. Note that outside of near-resonant states, tidal simulations are relatively insensitive to small-scale topographic changes and the blocking of the poles, so these measures have a negligible effect on the results (Wilmes & Green, 2014). See Fig. 1 for example bathymetries for four selected Cambrian time slices and present day.

**Fig. 1:**
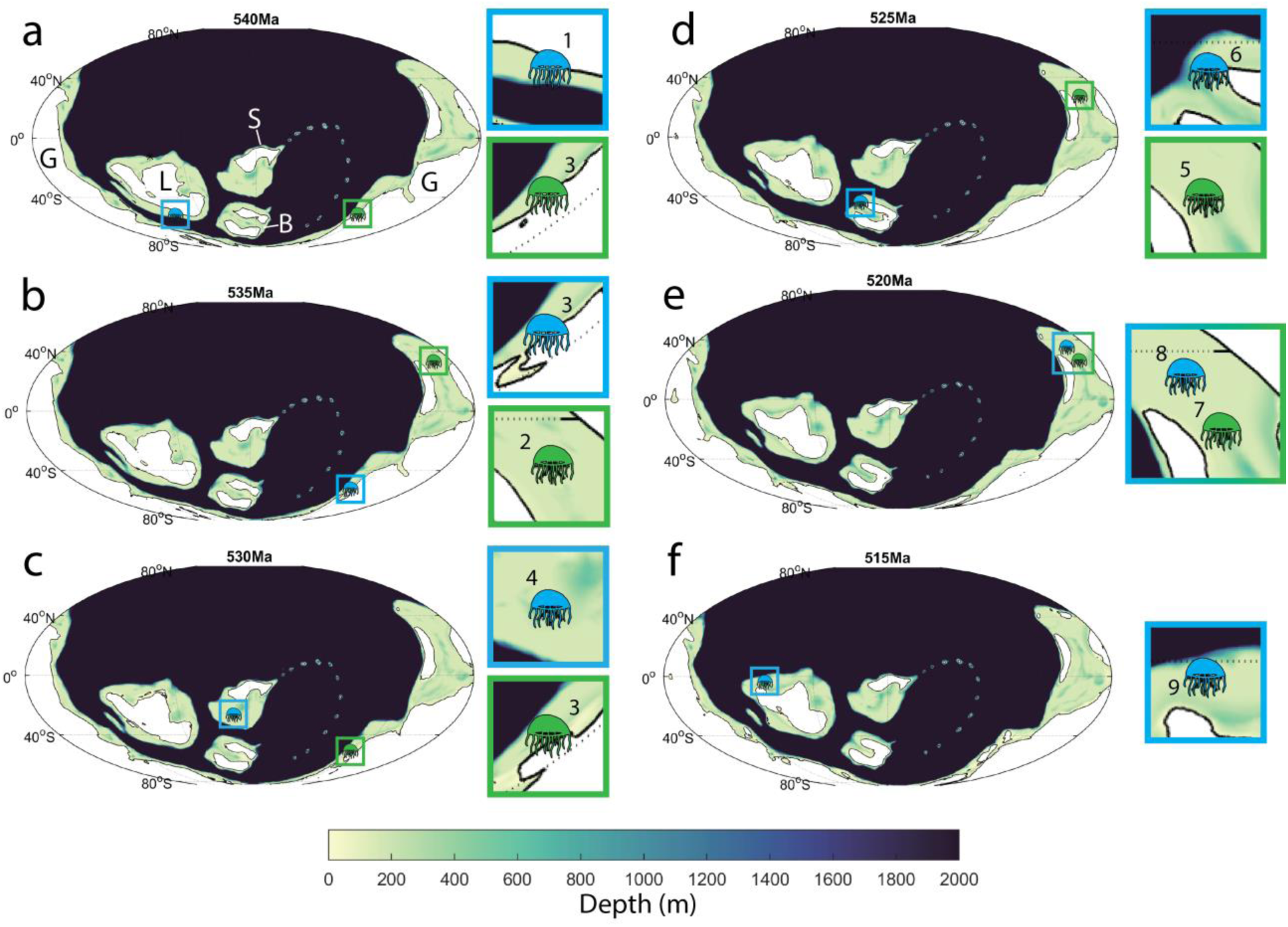
Bathymetry profiles for six time-slices at 5 Ma intervals from 540 – 515 Ma. Proxy locations for the relevant time slices are marked both on the full map and shown in more detail on the subplots to the right. Numbers in the subplots correspond to the proxy numbers in Table 1. Continents labelled in (a): B= Baltica, G=Gondwana, L=Laurentia, and S=Siberia

**Table 1:**
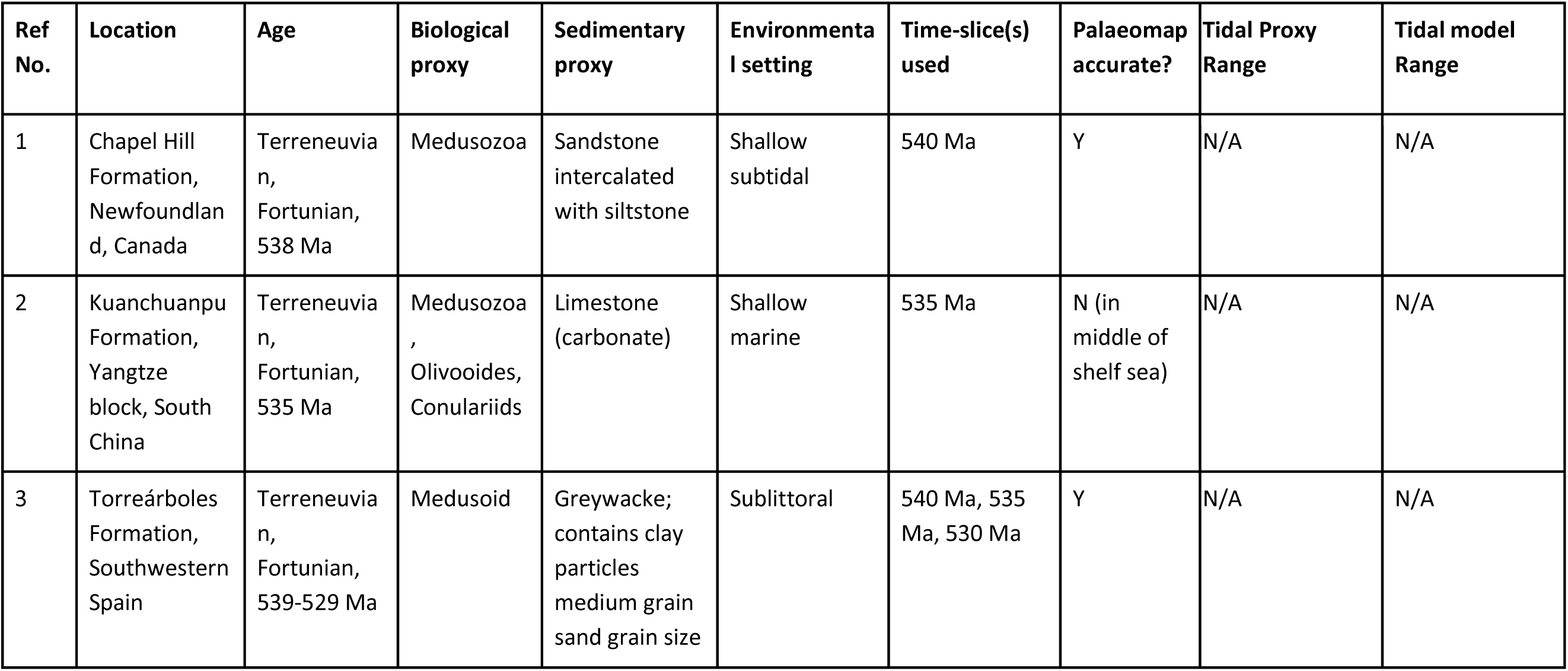
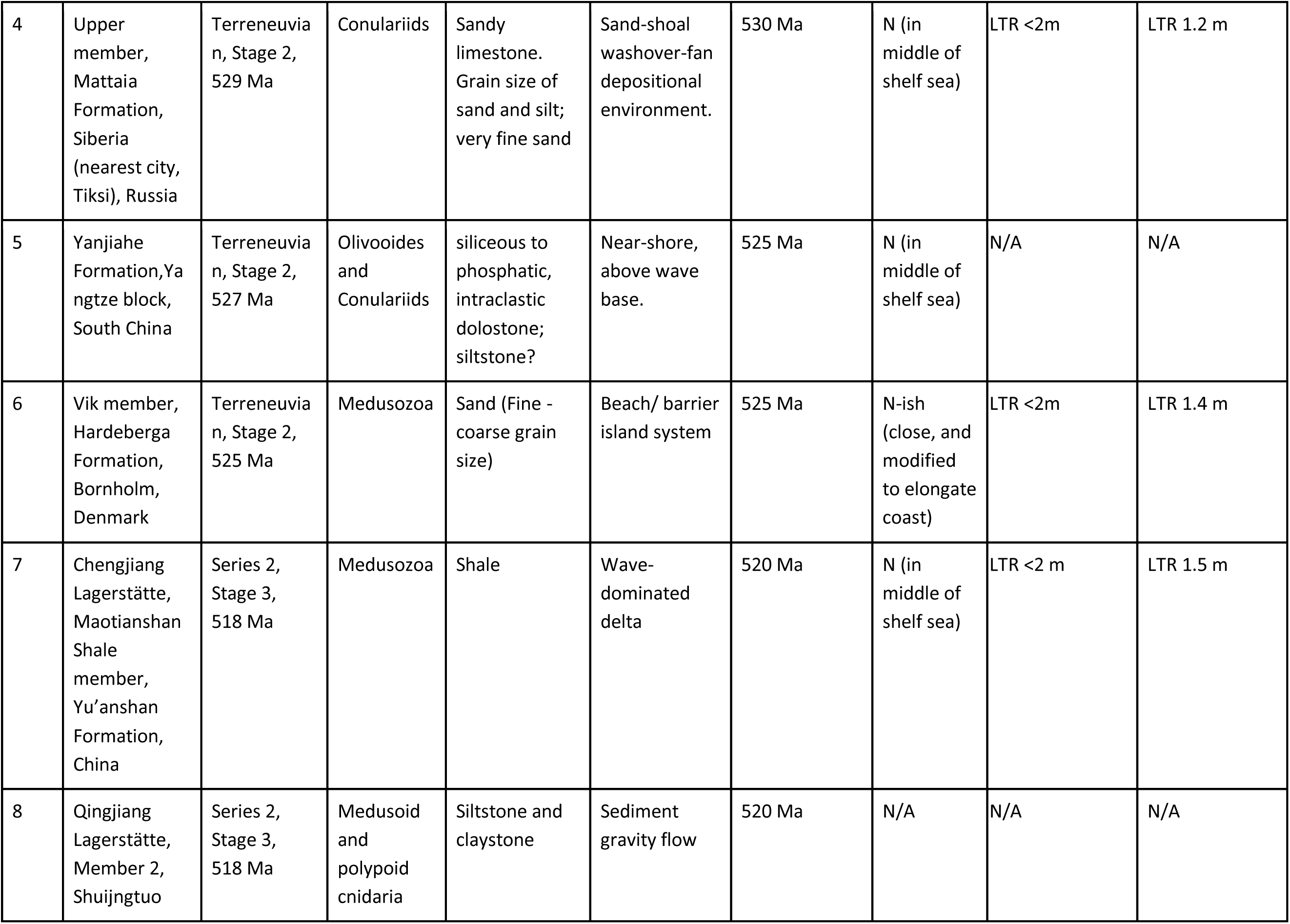
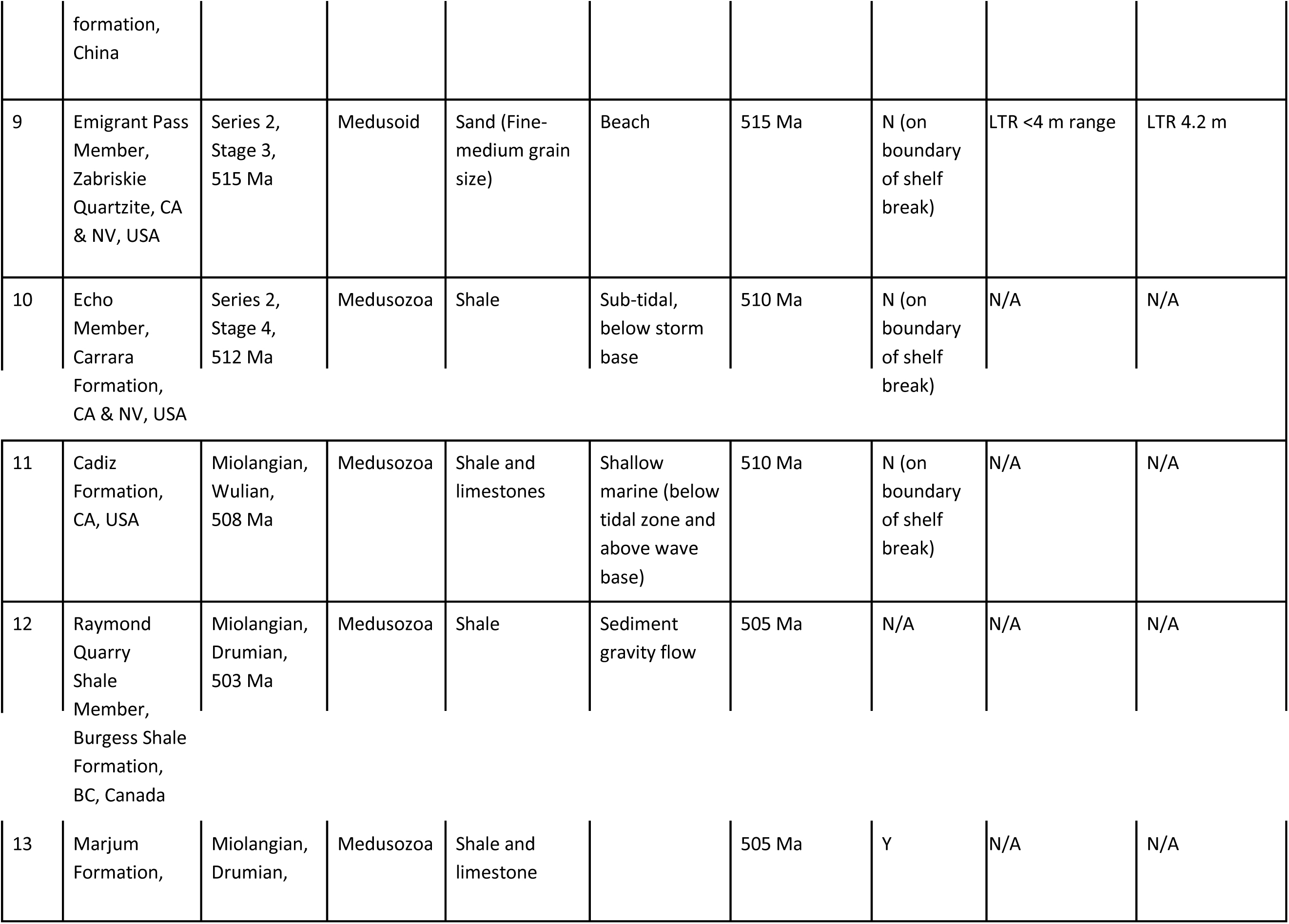
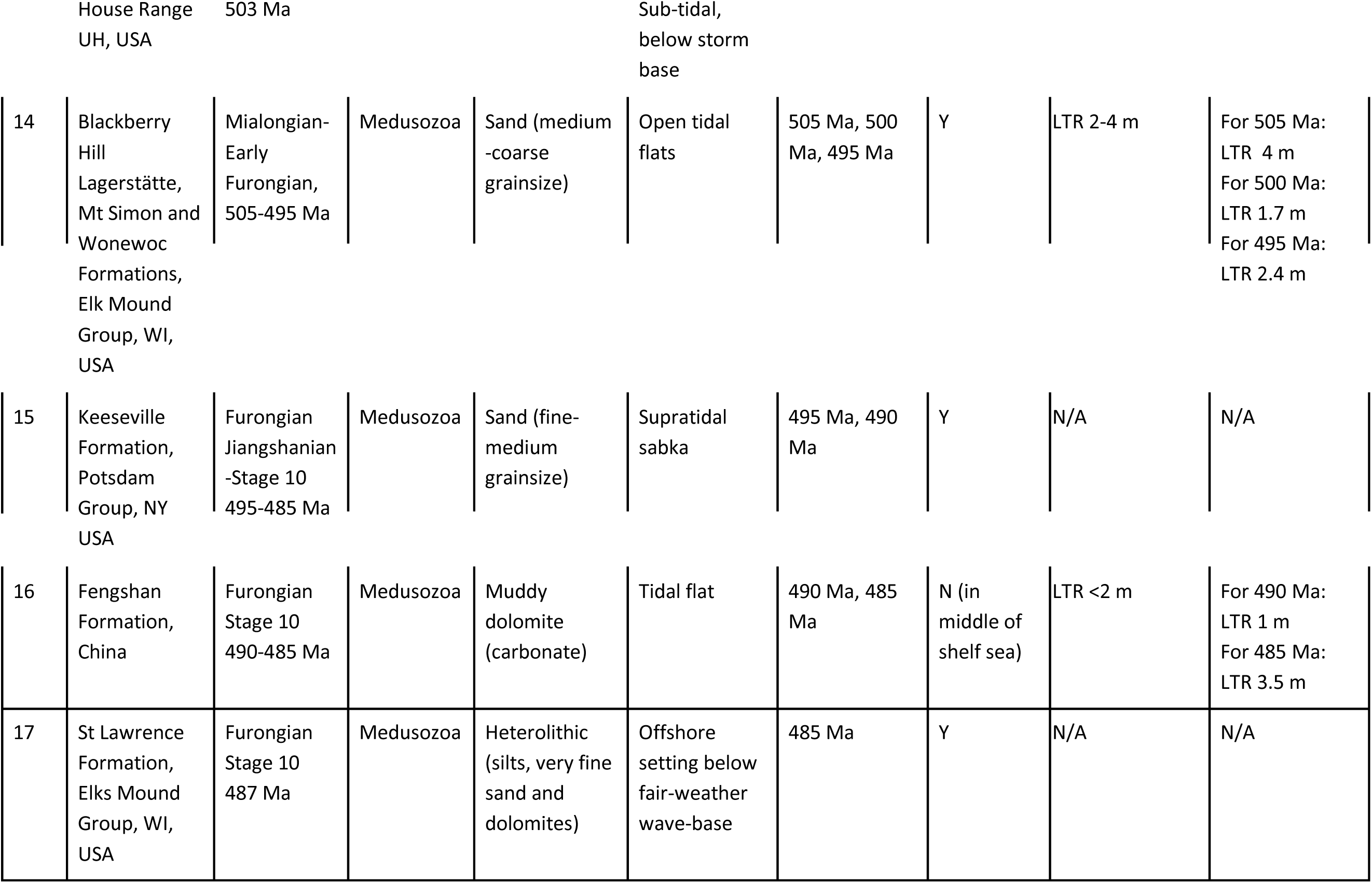
Sources and results summary. “Ref no.” is the reference number shown in Fig. 5a and related to the following references. 1: (Narbonne et al., 1991); **2**: (Dong et al., 2013; Wang et al., 2017; Wang et al., 2020); **3**: (Mayoral et al., 2021; Moreno-Martín et al., 2023); **4**: (Nagovitsin et al., 2015; Sarsembaev & Marusin, 2022); **5**: (Chang et al., 2020; J. Guo, Han, Iten, et al., 2020; J. Guo, Han, Van Iten, et al., 2020); **6**: (Clemmensen et al., 2017); **7**: (Han et al., 2016; Saleh et al., 2022; Young & Hagadorn, 2020); **8**: (Fu et al., 2019; Young & Hagadorn, 2020)**; 9**: (Barnes & Klein, 1975; Faggetter et al., 2017; Sappenfield et al., 2017); **10**: (Faggetter et al., 2017; Lieberman et al., 2017); **11**: (Waggoner & Collins, 1995); **12**: (Collom et al., 2009; Moon et al., 2023, p. 20; Young & Hagadorn, 2020); **13**: (Brett et al., 2009; Cartwright et al., 2007); **14**: (Braddy et al., 2022; Hagadorn et al., 2002); **15**: (Hagadorn & Belt, 2008; Lowe et al., 2017); **16**: (Latif et al., 2018; Song et al., 2021); **17**: (Hughes et al., 2000; Hughes & Hesselbo, 1997).

### 2.2 Numerical tidal modelling

The tides were simulated with OTIS – the Oregon State University Tidal Inversion Software - a numerical tidal model which has been used extensively to simulate deep-time, present-day, and future tides (Byrne et al., 2020; J. A. M. Green et al., 2020; J.A.M. Green et al., 2017; B. Guo et al., 2023, 2023; S.-B. Wilmes & Green, 2014) and it has been benchmarked against other forward tidal models and shown to perform well (Stammer et al., 2014). OTIS provides a numerical solution to the linearized shallow water equations,

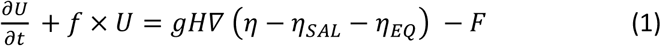

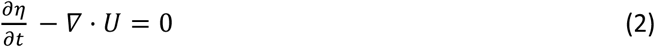

Here, **U**=**u***H* is the tidal volume transport (**u** is the horizontal velocity vector and *H* is the water depth), **f** is the Coriolis parameter, *g* is acceleration due to gravity, *η* is the sea-surface elevation, *η*_SAL_ is the self-attraction and loading elevation, *η*_EQ_ is the elevation of the equilibrium tide, and **F** the tidal energy dissipation term. The latter is a sum of two components, **F**_B_ and **F**_W_, where **F**_B_ parameterizes bed friction and **F**_W_ represents energy losses due to tidal conversion, i.e., the transfer of energy into a baroclinic tide. The first term is represented using the standard quadratic law, F_B_=*C*_D_ **u**|**u**|, where *C*_D_=0.003 is a dimensionless drag coefficient. The tidal conversion term is given by F_W_ = *C***U**, with a conversion coefficient, *C*, given by (J A M Green & Nycander, 2013; Zaron & Egbert, 2006)

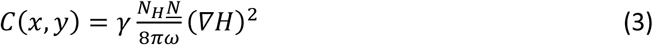

Here, γ = 50 represents a dimensionless scaling factor representing unresolved bathymetric roughness, *N_H_* is the buoyancy frequency at the seabed, *<u>N</u>* represents the vertical average of the buoyancy frequency, and ω is the frequency of the tidal constituent under consideration. The buoyancy frequency, *N*, is given by *N*^2^ = - g/ρ ∂ρ/∂z, where ρ is the density. Here, *N*^2^ was calculated from the density fields simulated HadCM3 and then used to compute globally averaged values of *<u>N</u>* and *N*_H_ for each time slice under investigation. The climate model (HadCM3BL-M2.1aD; Valdes et al., 2017) simulations are similar to the ‘Foster CO2’ (Foster et al., 2017) simulations of Valdes et al. (2021), except that they include improvements to some atmospheric and ocean parameters, and ocean physics. The simulations are identical to the ‘Scotese 07’ set of Judd et al. (2024), and are assigned ID ‘tfke’ in the “Providing Unified Model Access” (PUMA) database.

### 2.3 Simulations

We simulated the M_2_ (principal lunar), S_2_ (principal solar), and K_1_ (luni-solar declination) constituents for all time periods. In the following, we refer to the sum of the range of the three constituents as the Largest Tidal Range (LTR). Because of tidal friction, the Earth is spinning down and, to conserve angular momentum of the Earth-Moon system, the Moon is receding (Daher et al., 2021; MacDonald, 1964). Consequently, we changed the forcing parameters for our simulations and used a sidereal day length of 21.16 hrs, leading to a lunar period of 11.01 hrs and a lunar tidal forcing 11% above present-day. Note that the solar forcing has not changed since the Cambrian so the S_2_ forcing used was the same as for present day. Because ⅔ of the forcing for K_1_ comes from the Moon (and the remaining ⅓ from the Sun), the forcing for K_1_ was increased by 7.3%. The S_2_ and K_1_ periods were 10.60 and 21.16 hours, respectively.

### 2.4 Palaeoecology and proxies

Cnidaria are a major animal phylum comprising around 10,000 species, the majority of which are marine (Technau et al., 2015). The two major groups within Cnidaria are the Anthozoa, including sea anemones and corals, and the Medusozoa, including jellyfish. The primary differentiation of these groups is that Medusozoa possess a medusa stage in their life cycle, which the Anthozoa do not (Technau et al., 2015). Cnidaria first appeared in the fossil record during the late Ediacaran period (560-541 Ma), represented by medusozoans, and modern forms remain relatively similar to their ancient relatives (Van Iten et al., 2014).

There are numerous methods of soft tissue fossilisation, with one fossil capable of exhibiting several means of preservation. These include sediment casts or moulds, modification of organic material through diagenetic mineralisation (a process where soft tissue is replaced with minerals), and bioimmuration under multi-layered sheets of microorganisms, usually microbial mats (MacGabhann et al., 2019; Wilby et al., 1996; Young & Hagadorn, 2020). Here, medusae fossil deposits were identified through a literature review and from the Paleobiology Database (https://paleobiodb.org/). The majority of localities were identified in the fossil medusae review by (Young & Hagadorn, 2010), with a total of 17 useful cnidarian medusae localities found from all sources. The selection criteria consisted of the deposit having robust cnidarian medusae fossils and accompanying sedimentary data to inform on palaeoenvironment. The localities varied in age, spanning the entire Cambrian period, though several of the localities cluster at certain time intervals (see Table 1 for a summary). Localities that do not have an age that discreetly falls into one of the 5 Ma intervals have been represented across the multiple intervals within their age range.

We first assess whether the reconstructions are accurate for each of the proxy locations, because if the reconstructions are incorrect the tides will be too (and vice versa). The mapping was done through the GPlates online point reconstruction tool (https://gwsdoc.gplates.org/reconstruction/reconstruct-points) using present day latitude and longitude from the publications. To alleviate some of the uncertainty in the mapping, we present the model results as an average and standard deviation computed using a 5-by-5 gridpoint square centred around the proposed proxy location. In these calculations, land grid cells were ignored. Furthermore, there are two localities which are sediment gravity flow deposits, the Burgess Shale and Qingjiang (Collom et al., 2009; Fu et al., 2019). Because of this they do not accurately represent the environment that they are found in and will not be used to test reconstruction accuracy. Furthermore, in six of the locations (Mattaia Fm, Hardeberga Fm, Chengjiang lagerstätte, Zabriskie Quartzite. Blackberry Hill lagerstätte and Fengshan Fm, see Table 1 for details) the geological record was detailed enough to provide estimates of the tidal range at the time of deposition. This will allow us to make a direct comparison of the model outputs and the recorded ranges at these sites.

## 3 Results

### 3.1 Present-Day validation

The present-day control simulations from the 0 Ma bathymetry gives a root-mean-square difference of 11 cm for the M_2_ tidal amplitudes compared to the altimetry-constrained TPXO9 model (see Egbert & Erofeeva, 2002 and https://www.tpxo.net for details about TPXO, and Guo et al., 2023 and Wilmes et al., 2021 for modelling details). A simulation with a reduced-detail (degenerate) present-day bathymetry, with a resolution similar to reconstructed bathymetries from 50 Ma, produced a difference of about 20 cm (Green et al., 2017; Guo et al., 2023).

### 3.2 Proxies and reconstructions comparison

We will discuss the reconstructions in two parts: 540 – 515 Ma (Fig. 1) and 505 – 485 Ma (Fig. 2). In general, for 540 –515 Ma, the reconstructions fail to capture accurately the environments depicted in the proxies. The Chapel Hill Fm and Torreárboles Fm are accurately represented, they are both marginal marine from the sediment data and in the reconstructions (Fig. 1a-c). The biggest conflicts between sediment data and reconstructions are seen with the Mattaia Fm, Hardeberga Fm and Zabriskie Quartzite (Table 1; Fig. 1a-c,f). The Mattaia Fm and Zabriskie Quartzite represent sandy intertidal environments and are located far from shore in the reconstructions (Fig. 1c,f). The Hardeberga Fm too is located offshore in the reconstructions, however, the reconstruction was modified slightly to extend the nearby coastline and thus resulting in a more accurate marginal marine environment for the Hardeberga Fm (Fig. 1d).

**Fig. 2:**
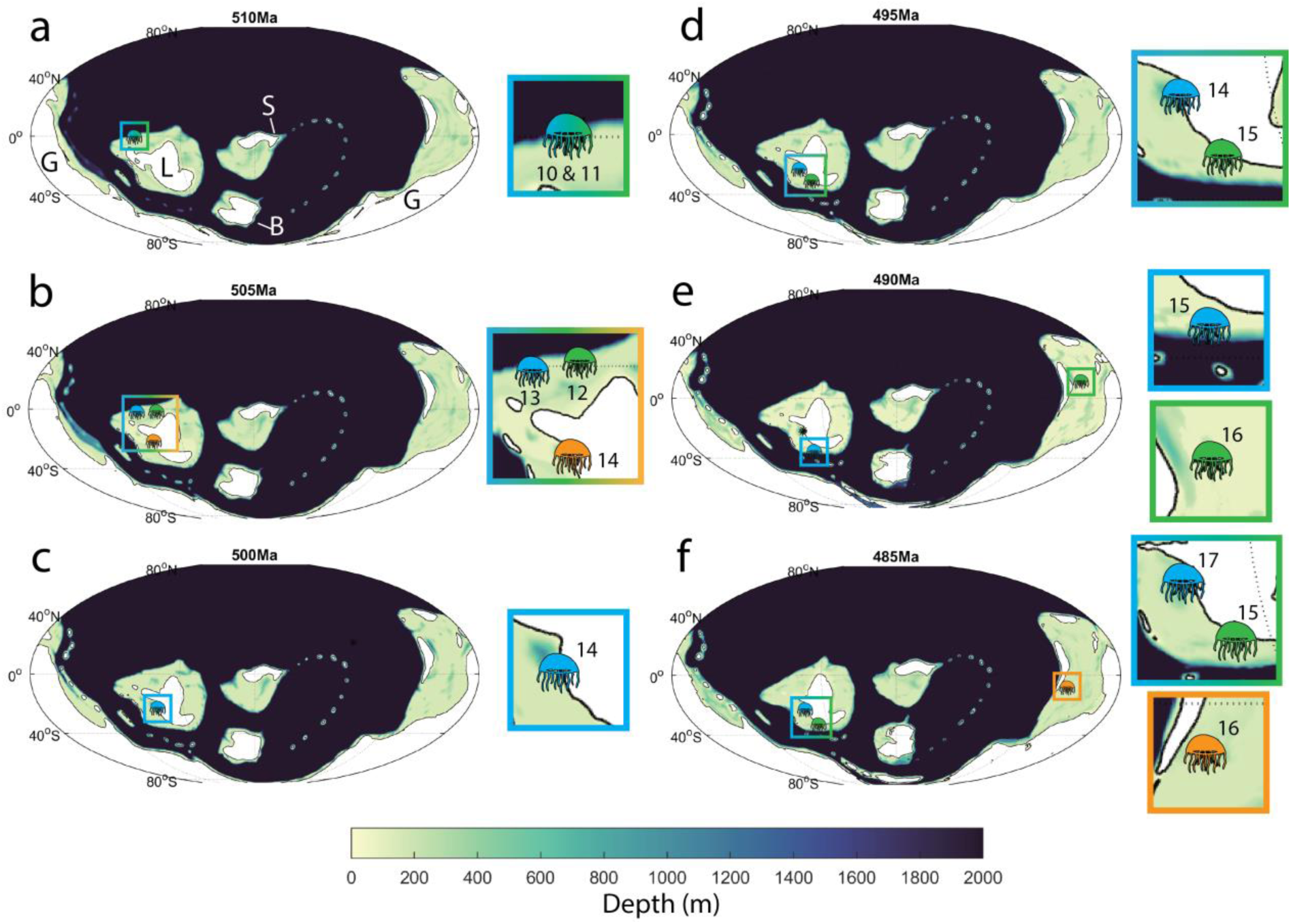
Bathymetry profiles for six time-slices at 5 Ma intervals from 510 – 485 Ma. Proxy locations for the relevant time slices are marked both on the full map and shown in more detail on the subplots to the right. Numbers in the subplots corresponds to the proxy numbers in Table 1. Continents labelled in (a): B=Baltica, G=Gondwana, L=Laurentia, and S=Siberia

For 510 – 485 Ma, there is an improvement in accuracy of correctly depicting the deposit environments in the reconstructions, but not for all (Table 1; Figs. 1 and 2). The Carrara Fm and Cadiz Fm represent sub-tidal deposits, making them difficult to constrain in which marine environment they are situated (Table 1). Nonetheless, they are situated far from a coastline and border the continental shelf break in the reconstructions, they were likely closer to the coastline (Fig. 2a and b). The remaining USA deposits (Blackberry Hill, Keeseville Fm and St Lawrence Fm) are accurately depicted in the reconstructions in relation to their paleoenvironments (Table 1; Fig. 2c-f). Deposits located in present-day China across the Cambrian are not accurately represented in the reconstructions across the entire Cambrian. These deposits are located in East Gondwana and are all marginal marine deposits, however, they are situated in shallow marine environments in the reconstructions, often far from a coastline (Figs. 1d,e and 2e,f).

### 3.3 Simulation results and comparison

We present tidal simulations for time-slices that possess at least one proxy with tidal range information (Table 1). The Largest Tidal Range (LTR) is shown, with a global overview and a zoomed-in view for the location of each proxy. Note that in the following, we define tidal ranges as microtidal (range <2 m), mesotidal (2-4 m range), and macrotidal (4-8 m range).

#### 3.3.1 530 Ma

There are several localised meso-macro tidal regions in the simulation for 530 Ma around each of the continents, with greatest abundance in East Gondwana (Fig 3a). The Mattaia Fm tidal proxy is situated SW of Siberia, and in our model exhibits an average LTR range of 1.2m (Table 1; Fig. 3b). This model output matches with microtidal range estimate from the proxy (Table 1).

**Fig. 3:**
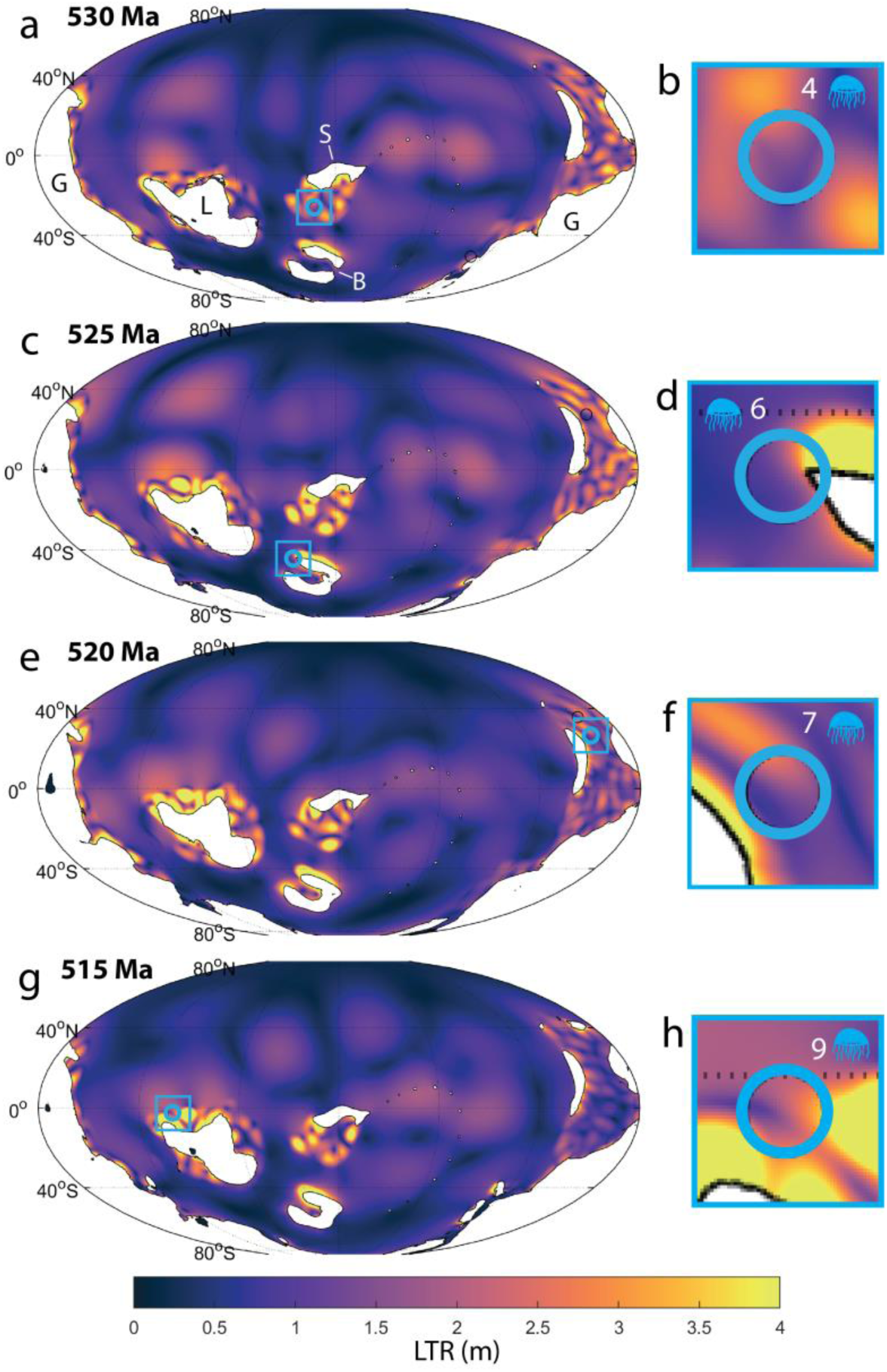
Tidal simulation outputs showing Largest tidal range (LTR, in metres). Global reconstructions are shown for (a) 530 Ma, (c) 525 Ma, (e) 525 Ma and (g) 515 Ma, with corresponding magnified views of tidal proxy palaeolocations provided in panels (b) (Mattaia Fm), (d) (Hardeberga Fm), (f) (Chengjiang Lagerstätte), and (h) (Zabriskie Quartzite). Continents labelled in (a): B=Baltica, G=Gondwana, L=Laurentia, and S=Siberia

#### 3.3.2 525 Ma

Relative to 530 Ma, tides are noticeably more energetic, with prominent macrotidal regions present north of Laurentia and Baltica and along the southern Siberian coast (Fig. 3a,c). The Hardeberga Fm proxy is located NW of Baltica and represents a barrier island system, which is associated with microtidal ranges. In our simulation the average LTR is 1.4m, which fits with the proxy estimate (Table 1; Fig. 3d).

#### 3.3.3 520 Ma

Generally, the global tidal pattern is similar between 525 Ma and 520 Ma, with a notable decrease in tidal ranges around East Gondawana (Fig. 3c,e). Our tidal proxy, Chengjiang lagerstätte, is located in East Gondwana, where the tides have become less energetic compared with 530 Ma. From the model, the proxy has a LTR range of 1.5m, which matches with the microtidal regime depicted by the sediment data (Table 1; Fig. 3f).

#### 3.3.4 515 Ma

The LTR has continued to decrease in East Gondwana, with mesotidal regions becoming more localised. Elsewhere, macrotidal regions are still seen in Laurentia, Siberia and Baltica but have become reduced in extent (Fig. 3g). The Zabriskie Quartzite proxy is situated NW of Laurentia, and lies on the edge of a macrotidal regions, and has an average LTR of 4.2m (Table 1; Fig. 3h). This is slightly higher than the range suggested by the proxy, of a mesotidal range (Table 1).

#### 3.3.5 505 Ma

The global tidal pattern broadly mirrors that of 515 Ma, with the notable exception of an amplification of tides within the seaway off East Gondwana, where several macrotidal regions emerge (Figs. 3g, 4a). The Blackberry Hill Lagerstätte tidal proxy is located on the west coast of Laurentia. In our simulation, this locality is situated within a macrotidal regime with an average LTR of 4.8m, which exceeds the 2–4 m tidal range depicted by the proxy data (Table 1; Fig 4b).

**Fig. 4:**
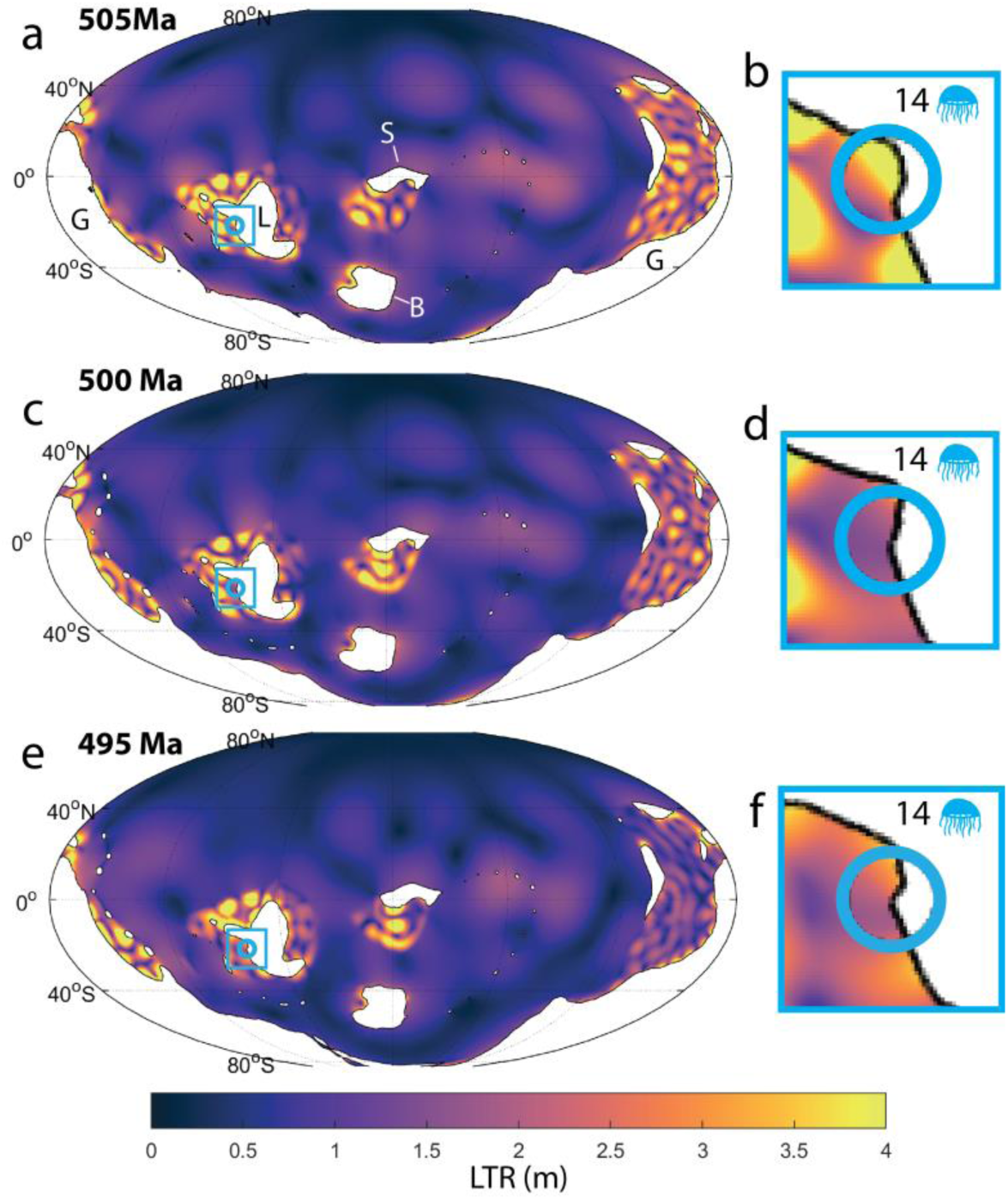
Tidal simulation outputs showing Largest tidal range (LTR, in metres). Global reconstructions are shown for (a) 505 Ma, (c) 500 Ma, and (e) 495 Ma, with magnified views of the tidal proxy palaeolocation for the Blackberry Hill Lagerstätte tidal proxy (b,d,f). Continents labelled in (a): B=Baltica, G=Gondwana, L=Laurentia, and S=Siberia

#### 3.3.6 500 Ma

At 500 Ma, macrotidal regions around Laurentia and Siberia remain similar to those at 505 Ma, with an expansion of macrotidal zones around West Gondwana and a corresponding decrease in East Gondwana (Fig. 4c). Notably, macrotidal regimes are absent around Baltica. In our simulation, the Blackberry Hill Lagerstätte lies within a region featuring an average LTR of 2 m, which at the lower end of the proxy-estimated mesotidal range of 2–4 m (Table 1; Fig. 4d).

#### 3.3.7 495 Ma

There are broad similarities in the tidal patterns of 500 Ma and 495 Ma, though tidal ranges decrease in the seaway in East Gondwana for the latter interval (Fig. 4e). In our simulation, the Blackberry Hill Lagerstätte lies in a meso-tidal region with an average LTR of 3.4m this aligns well with the proxy estimate of a mesotidal regime (Table 1; Fig. 4f).

#### 3.3.8 490 Ma

Between 495 Ma and 490 Ma, a notable decrease in tidal ranges occurs around West and East Gondwana, while the distribution of macrotidal regions remains stable around Laurentia, Siberia, and Baltica (Figs. 4e and 5a). The Fengshan Formation is located in East Gondwana, southeast of an elongated island (Fig. 5a). In our simulation, the LTR at this proxy locality is 2 m, which aligns well with the proxy-estimate of a micro- to mesotidal regime (Table 1; Fig. 5b).

**Fig. 5:**
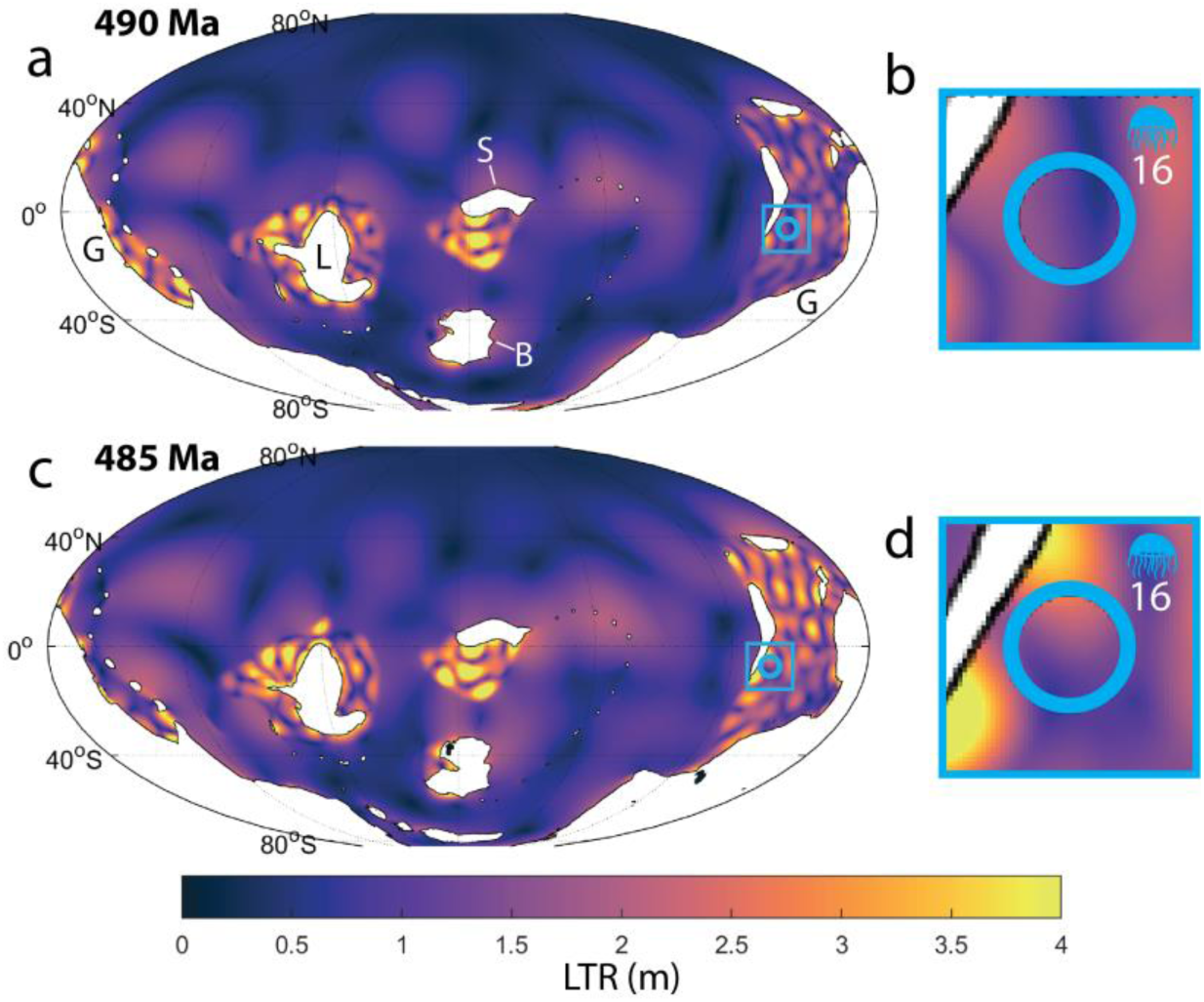
Tidal simulation outputs showing Largest tidal range (LTR, in metres). Global reconstructions are shown for (a) 490 Ma and (c) 485 Ma, with magnified views of the tidal proxy palaeolocation for the Fengshan Fm tidal proxy (b,d). Continents labelled in (a) : B=Baltica, G=Gondwana, L=Laurentia, and S=Siberia

#### 3.3.9 485 Ma

At 485 Ma, regions featuring meso- to macrotidal ranges reemerge in East Gondwana, accompanied by an expansion of macrotidal regions around Laurentia, Siberia, and Baltica, and a concurrent decline in tidal ranges across West Gondwana (Fig. 5a,c). In our simulation, The Fengshan Fm is situated in an area with an average LTR of 3.4m which exceeds with the proxy-estimate of ∼2m (Table 1; Fig. 5d).

### 3.4 Synthesis

The tidal range estimates have been collated along with estimates from the model and synthesised in Fig. 6. For several locations, only an age range had to be considered, so these locations are presented in time-slices that occurred within their respective age ranges in panels c) and d) of Fig. 6. The simulated LTR (Fig. 3-5) match the proxies well. Zabriskie Quartzite at 515 Ma is deemed to have been mesotidal (<4m range) and the model sits just above that at 4.2 m, whereas for Blackberry Hill at 500 Ma the model again sits just under the 2-4 m range predicted by the proxies. Some of the formations are poorly constrained in time, so in Fig. 6d we present results for the median time slices for these locations. The LTR from Blackberry Hill (Table. 1;Fig. 5, data at 505-495 Ma), shows that the model works well for 505 and 495 Ma (with 495 Ma better than 505 Ma) but less so at 500 Ma. Consequently, the correct dating is important as well and it is a tantalising result that maybe this type of work can also help to constrain the age of the outcrops.

**Fig. 6.**
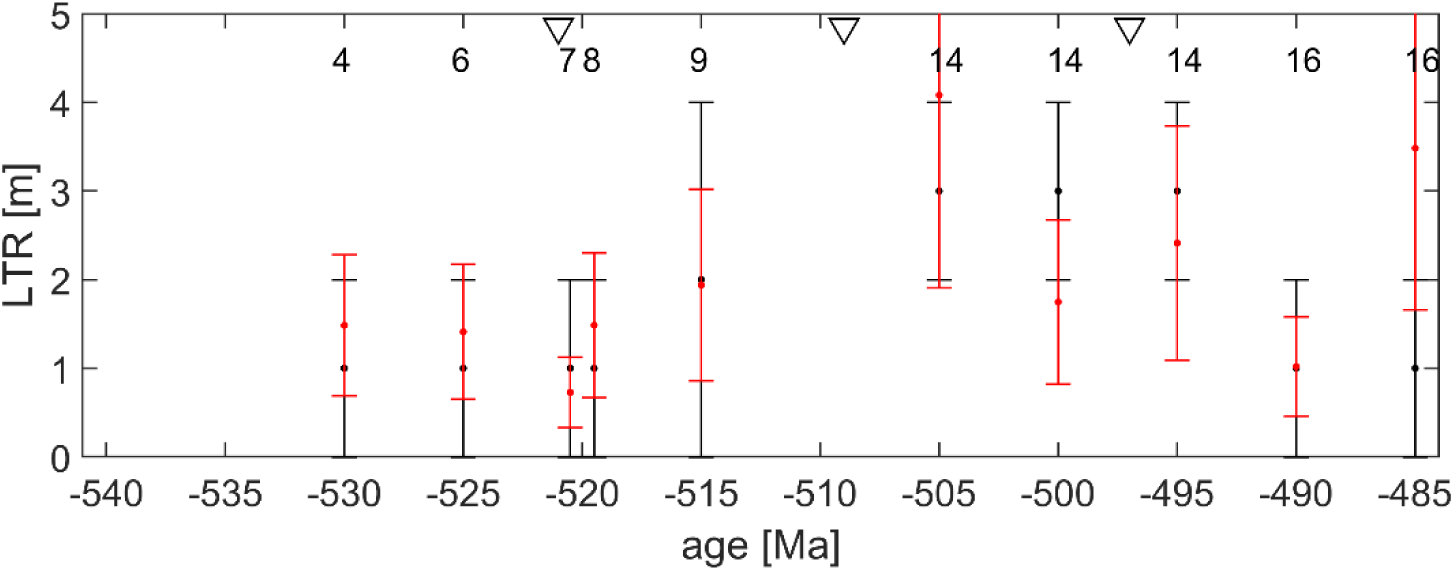
Comparison between the model results (red) and proxies (black) for Largest Tidal Range (LTR). Note that at time slices with more than one location, the data is plotted with a 0.5 Ma offset in age for clarity in all panels. The numbers at the top of the panel show the reference number from Table 1. The proxies are plotted as smallest to largest tidal range in the whiskers with the dot marking the average range. The model results are the mean ± standard deviation of the model output within the 2-by-2 gridpoint square centred over the proxy location (see methods for details). Note that references 14 and 16 are poorly constrained in time, so have been repeated over the time slices they cover; see the text for details. Furthermore, note that location 14 for 505 Ma goes off scale in the model data.

## 4 Discussion

There is mixed success when it comes to mapping the proxies on the reconstructions. This includes a persistent issue with the location of the Cambrian proxies from China in the reconstructions: they all should be located near a shoreline. We are confident that such small changes to the topography would not greatly change the tidal signal in the model simulations. For the formations where the reconstructions and proxy environment disagree, i. e., the locations in present day China, Zabriskie Quartzite, Carrara Fm and Cadiz Fm in the US, and the Mattaia Fm (Table 1; Figures 1,2), larger changes to the reconstructions may be necessary. This is unless these locations are located close to islands which may not be resolved at the 1/10° resolution in the reconstructions we use. A small amendment to the reconstruction was made for the case of the Hardeberga Fm, and this resulted in a more accurate representation both in the reconstruction and tidal simulation. Our study demonstrates the importance in consulting the fossil record and respective paleoenvironments in producing accurate paleogeographic reconstructions, in line with other studies (Cao et al., 2017; Kocsis & Scotese, 2021).

The tidal simulations highlight the spatial variability of tidal ranges on local, regional, and global scales. This is illustrated by the mosaic of hotspots with large tidal ranges in the seaway of East Gondwana throughout all the simulations (Figs. 4-6). Similar signals are found around Laurentia and off the south coast of Siberia, with a variation in tidal ranges around Baltica. These hotspots all occur in shelf seas and may be caused by an enhanced tidal amplitude because of the shallow water, or because of local tidal resonances. It highlights that one location cannot be used as a proxy for another location.

We argue that the use of fossils, in this case of cnidarian medusae, has proven effective to obtain paleoenvironmental information for a variety of marine settings. Fossil localities were found at regular intervals spanning the entire Cambrian, providing temporal coverage. They also covered most of the large continents present during the Cambrian, providing a relatively good spatial representation. The diverse habitats that are represented by the fossil localities is of great benefit for constraining reconstructions, particularly coastal bathymetry. This is less beneficial for constraining tidal simulations; proxies that lie outside of the intertidal zone cannot provide information on tidal range estimates. Nonetheless, 6 of the 17 proxies yield tidal range estimates.

Tidal ranges provide the most accurate tidal model results. It has been documented in other deep-time tides studies (e.g., Byrne et al., 2020; Mángano et al., 2014) that tidal range estimates from proxies are robust, and we argue that the overall agreement between our tidal range results and the proxies can be trusted. It is also encouraging that the current speed estimates match closely with the simulated tidal currents, and we are confident that our tidal model results are as reliable as they can be for simulations of a period 540-485 Ma.

The Cambrian period also marks another significant milestone in the evolution of life as it marks the first evidence of animals moving on to land, in the form of tracks from soft-bodied organisms over tidal flats. The intertidal zone is home to members from all major animal phyla, making it a key environment despite its relatively small size globally. This environment is critically significant during the Cambrian period as all animals are marine during this time and there is an observed shift in faunal dominance from offshore into marginal-marine ecosystems (Wood et al., 2019). It has been proposed that the radiation of animals from the ocean to land occurred because animals were stranded during high tides (Braddy et al., 2022; Collette et al., 2010; Young & Hagadorn, 2010, 2020), and the results from our study support this idea. Some of the oldest signs of life on land come from Blackberry Hill in present day Wisconsin, USA, which is part of the Mt Simon and Wonewoc Formations (our location 14; see also Braddy et al., 2022 and Hagadorn et al., 2002) where we find a mesotidal range in both proxies and model (Fig. 6d). Tidal influence has also been proposed as a driver in the water to land transition of arthropods and vertebrates later in the Palaeozoic (Byrne et al., 2020; Mángano et al., 2014).

The Cambrian saw a significant evolution in abiotic factors in our oceans, establishing those we see at Present Day. During the Precambrian and into the Cambrian, ferruginous and euxinic conditions were dominant resulting in a dynamic chemocline and oxygen minimum zones were abundant. These redox conditions would have restricted life-forms to be small in size. Although these conditions were mainly confined to deeper water environments, the chemocline frequently shoaled into shallower waters during periods of transgression (Wood et al., 2019). The unstable redox conditions as a result likely caused high faunal overturning and could have acted as an evolutionary driver (Wood et al., 2019; Wood & Erwin, 2018). However, the spatio-temporal details on the evolution of redox conditions from the Precambrian into the Ordovician are poorly understood. Recent efforts on a holistic integration of various parameters including palaeontology, geochemistry, and modelling have been deployed to achieve a greater understanding (Wang et al., 2018; Wood et al., 2019). The results from our tidal model could provide further information on this topic. A shoaling of the chemocline points to a stronger vertical water column stratification, or a stratification which became established, due to the transgression. This would happen because as the water column deepens, the tides can no longer fully mix it and a (seasonal) stratification can become established (Simpson & Bowers, 1981; Simpson & Hunter, 1974). Tides would therefore play a critical role in the regulation of redox conditions in marginal marine environments, and further high-resolution work into tide, both through proxies and the model, during the Cambrian is warranted.

We have demonstrated that the use of fossils and their respective paleoenvironments is a viable way to constrain both palaeo-tidal model simulations and the palaeogeographical reconstructions. The technique can be applied to other time periods using different coastal marine taxa; note that the cnidarian medusae preservation window was optimal during Cambrian and are less abundant after this period (Young & Hagadorn, 2020). Important evolutionary events where marginal-marine environments play a key role should be of high priority to investigate, and has been proven successful in a study on tides during the fish to tetrapod transition (Byrne et al., 2020). Notable events to explore could be the water-to-land transition of arthropods and the near-shore cradle origin for vertebrates during the Ordovician and Silurian, respectively (Mángano et al., 2014; Sallan et al., 2018). Extinction events which impact marginal-marine environments are also worth studying from a tidal perspective, e.g. Kellwasser extinction event during the Devonian. This study echoes the importance and need for holistic and interdisciplinary research to gain a more accurate depiction of the deep-time Earth system.

## CRediT authorship contribution statement

### Author contribution

The idea came from JAMG, who along with IP did the simulations using CRS’ reconstructions and PJV’s/DJL’s stratification data. HAMB and MEHH collated the proxy information and did the analysis. All authors contributed to the writing led by HAMB.

## Acknowledgements

The research was funded by the UK Natural Environment Research Council through MATCH (NE/S009566/1; awarded to JAMG, with support for IP, HAMB, and MEEH). HAMB also acknowledges current funding under the European Union’s Horizon Europe research and innovation program by the Marie Skłodowska-Curie grant agreement No.101150146 (MOSAIC). MEHH also acknowledges a Bangor University Summer Internship and a Santander Undergraduate Scholarship. The tidal model simulations were done on Supercomputing Wales, which is part-funded by the European Regional Development Fund (ERDF) via the Welsh Government. DJL and PJV acknowledge NERC grant PaleoGradPhan (NE/X000222/1).

## Data availability

The reconstructions can be found on https://www.earthbyte.org/paleodem-resource-scotese-and-wright-2018/. The tidal model simulation results will be available at Zenodo, doi: 10.5281/zenodo.10819584 after publication and can be requested for review. The tidal model code is available from the corresponding author upon request or from https://www.tpxo.net/otis.

